# Adaptive evolution of *P. aeruginosa* in human airways shows phenotypic convergence despite diverse patterns of genomic changes

**DOI:** 10.1101/2023.07.19.549629

**Authors:** Akbar Espaillat, Claudia Antonella Colque, Daniela Rago, Ruggero La Rosa, Søren Molin, Helle Krogh Johansen

## Abstract

Selective forces in the environment drive bacterial adaptation to novel niches, choosing the fitter variants in the population. However, in dynamic and changing environments, the evolutionary processes controlling bacterial adaptation are difficult to monitor. Here, we follow 9 cystic fibrosis patients chronically infected with *Pseudomonas aeruginosa*, as a proxy for bacterial adaptation. We identify and describe the bacterial changes and evolution occurring between 15 and 35 years of within host evolution. We combine whole genome sequencing, RNAseq and metabolomics, and compare the evolutionary trajectories directed by the adaptation of four different *P. aeruginosa* lineages to the lung. Our data suggest divergent evolution at the genomic level for most of the genes, with signs of convergent evolution with respect to acquisition of mutations in regulatory genes, which drive the transcriptional and metabolomic program at late time of evolution. Metabolomics further confirmed convergent adaptive phenotypic evolution as documented by reduction of the quorum sensing molecules acyl-homoserine lactone, phenazines and rhamnolipids (except for quinolones). The modulation of the quorum sensing repertoire suggests that similar selective forces characterize at late times of evolution independent of the patient. Collectively, our data suggest that similar environments and similar *P. aeruginosa* populations in the patients at prolonged time of infection are associated with an overall reduction of virulence-associated features and phenotypic convergence.

**Summary:** Selective forces in the human environment drive bacterial adaptation to novel niches, choosing the fitter variants in the population. We have investigated the evolutionary processes in 9 CF patients infected with *Pseudomonas aeruginosa* in the airways for several decades. To describe the within host evolution and trajectories of four different *P. aeruginosa* lineages to the lung environment we have combined whole genome sequencing, RNAseq and metabolomics. In this patient cohorte with persistent bacterial infections our data suggest that similar environments and similar *P. aeruginosa* populations in the patients at prolonged time of infection are associated with an overall reduction of virulence-associated features and phenotypic convergence

## Introduction

Microbial adaptation to a particular environment is directed by specific selective forces, and each niche represent a unique fitness landscape for the infecting bacteria. In this scenario, the selection of beneficial mutations that fix and expand in the population help bacteria to successfully adapt and persist. Yet, monitoring the occurring adaptational events in dynamic natural environments, and inferring the driving selective pressures, remains a challenge, due to i) complex spatial-temporal fluctuating conditions (e.g., temperature, pH, osmolality, nutrient gradients), ii) population dynamics (e.g., prey-predator, mutualistic relationship, or pathogenic interactions), and iii) interkingdom-interactions (host-microbe interactions). Genetic variations in such bacterial populations have, therefore, been difficult to associate with specific adaptive processes, if the complex conditions are at least not transiently stable. Consequently, mutation acquisition as a proxy for the selective pressures is usually insufficient to validate the particular significance of the specific genetic modifications (1).

We investigate bacterial adaptation and evolution of *Pseudomonas aeruginosa* (Pa) in the airways of people with cystic fibrosis (pwCF) during the progression of colonization and infection. The CF lung infection model offers unique opportunities, as there is extensive within-patient follow-up information, CF sputum samples are routinely sampled from patient cohorts to diagnose bacterial infection status, and detailed characterization of CF lung disease progression has been well documented (2, 3). Considering the complexity and dynamics of the human airways, including the particular multi-species microbial communities described in CF airways, we suggest that our findings concerning bacterial adaptation in this environment reflect evolutionary processes occurring generally in many other natural environments (4, 5).

Previously, the evolutionary dynamics of a persistent and highly successful Pa lineage, DK02, that had disseminated to more than 40 patients in the Copenhagen CF clinic, has been described (4), with respect to the genomic and phenotypic changes over a period of more than 200,000 bacterial generations (4). The DK02 lineage shows limited inter-patient diversification, after an initial period of rapid adaptation. This is most likely caused by acquisition of a few regulatory mutations affecting the transcriptional profile followed by a period of genetic drift with minor transcriptional changes (4, 5).

This opened the question of whether the DK02 evolutionary history could be used as a reference to predict the adaptive pathways for other Pa lineages, when adapting towards a state of chronic infections in human airways. To address this question, we here investigate the evolution of three alternative and wide-spread persistent Pa lineages, DK01 (6), DK19 (PA14) (7, 8) and DK06 (C-clone) (9, 10), in comparison with the DK02 evolutionary pathway (3). Specifically, we resolved the evolutionary history of *P. aeruginosa* in the airways of several chronically infected pwCF, as they reflect an adapted population to the lung environment (3). The strategy has been to correlate the acquired mutations identified in the respective genomes with the globally expressed transcriptional network in DK02, and the resulting biosynthetic products. Overall, we document the value of using combinations of omics approaches to better understand evolutionary dynamics in complex environments.

## Results

### Within-patient genome evolution: Divergence and convergence of Pa lineages

To characterize the evolutionary trajectories securing the persistence of Pa in pwCF, we investigated a collection of Pa clinical isolates sampled longitudinally between 1973 and 2021 from 9 patients attending the Copenhagen CF Clinic (Table S1). For each patient, we compared initial isolates (referred to as “early”) collected within 2 years of the diagnosis of chronic infection, with isolates collected after 15 years (DK06 and DK19 lineages), and 35 years (DK02 lineage) (referred to as “late”) of infection (Fig.1A, Table S1). For the DK01 lineage, no early isolates were available in our collection, and therefore, “intermediate” strains isolated after more than 10 years after the onset of chronicity and evolved for 35 years in each patient were used (Fig.1A, Table S1). This collection comprises isolates with evolutionary histories estimated to cover between 35,000 and 150,000 bacterial generations (Table S2).

**Figure 1.**
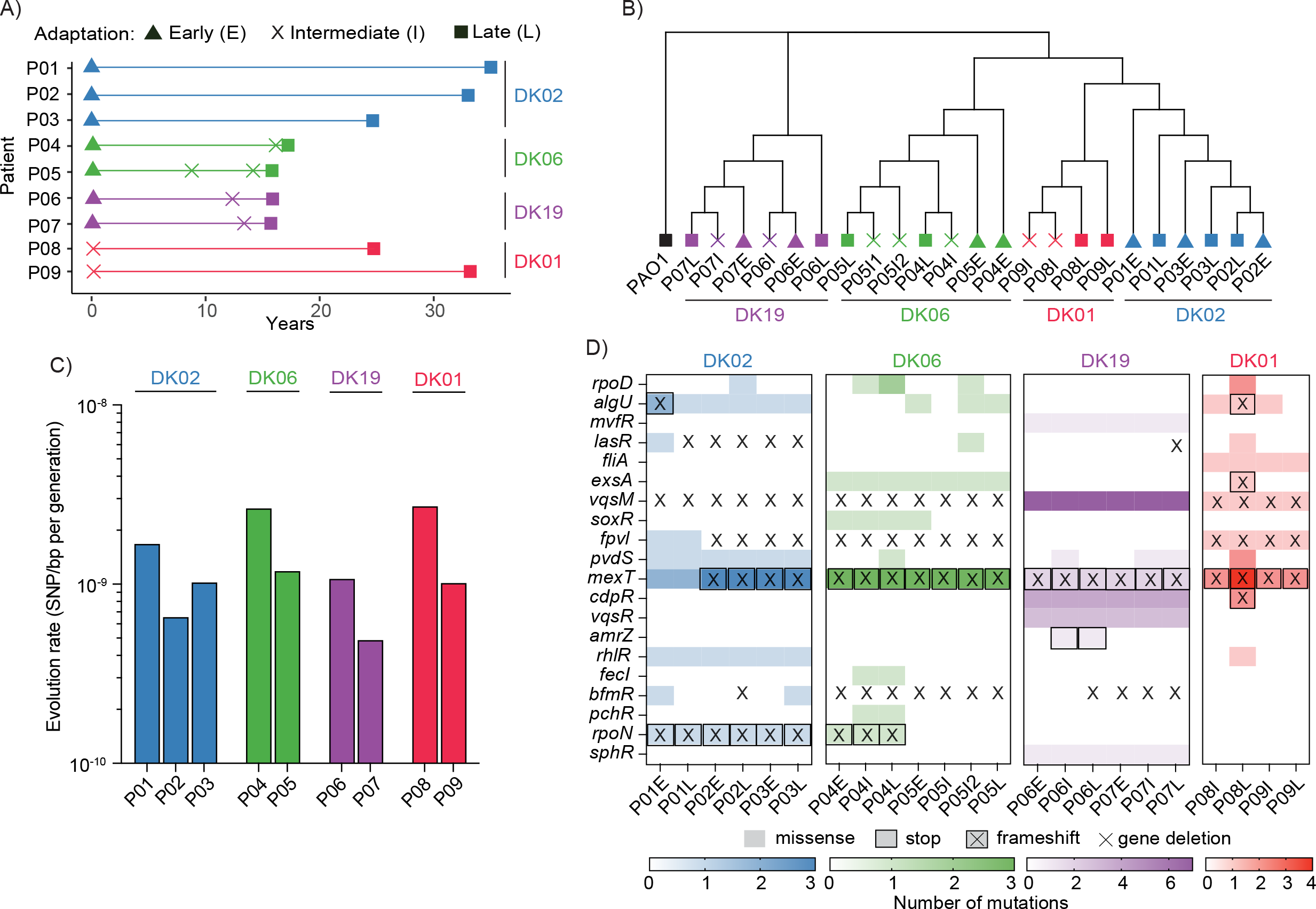
Distribution of the lineage-associated mutations. A) Longitudinal isolated samples from cystic fibrosis patients. Lineages dominating each patient are color-coded (DK01 red, DK02 blue, DK06, green and DK19 purple). B) Evolutionary history of the isolates represented by Maximum likelihood reconstruction. The bootstrap (500 replicates) tree is based on the concatenated SNPs of each isolate relative to PAO1. C) Estimation of evolution rates as SNPs per bp per generation time. D) Heatmap of the mutations found within regulatory genes.

Single nucleotide polymorphisms (SNPs) were used as phylogenetic markers to reconstruct the evolutionary history of the strains. As previously observed (11), the genomes grouped primarily according to their lineage, suggesting a strong evolutionary contingency (Fig. 1B). Since niche specialization depends on the constant modulation of the mutation rate (12), we evaluated if the mutation rates in the lineages differed. Despite significant differences in numbers of generations, neither the synonymous nor the non-synonymous mutation rates were found to differ between lineages (Fig. 1C).

When comparing the introduced genetic changes across lineages, we found evidence for both convergent and divergent evolution within and between lineages. Among the mutated genes identified in the early/intermediate isolates, we found ∼31% of these to be shared across the samples (Figure S1A). Specifically, all lineages showed convergent acquisition of mutations in genes related to i) virulence pathways (e.g., Type II and III Secretion apparatus, iron homeostasis, ii) antibiotic resistance (e.g., efflux pumps systems), and iii) motility (Table S3). These traits are known to be frequently lost after establishment of chronic *P. aeruginosa* infections in CF airways (1). When comparing the late isolates, ∼34% of the mutated genes were shared (Figure S1B, Table S3). All late isolates shared i) indel mutations in the *tonB1* transporter, a protein that is required for the uptake of ferripyoverdine (13), ii) the same missense mutation in the iron-translocating oxidoreductase *rnfC*, and iii) mutations in two component of the Type VI Secretion system (Table S3). Among the accumulated mutations from early/intermediate to late isolates, only low percentages (∼3% for DK01 and DK02, 10% for DK06 and 18% for DK19) of mutated genes were shared within each specific lineage. In contrast, similar categories of genes and biological pathways were shared across the different lineages (Figure S1C and D). Overall, our data suggest that early mutational patterns show partial convergence at the genomic level, whereas at later stages of infection there is a strong within-patient specialization of the particular isolate/lineage (genomic divergence).

The repeated occurrence of pathoadaptive mutations suggests convergent mechanisms of adaptation at both the genomic and the phenotypic level (4, 11, 14). Interestingly, regulatory mutations show evolutionary convergence, either in a lineage-independent manner (fixed in all lineages), or with reference to time (early/intermediate ➔ late), patient or lineage (Fig.1D and Table S4). In the case of the lineage-independent mutations, missense and/or frameshift mutations were identified in the genes encoding the multidrug efflux pump regulator MexT and in the virulence modulator VqsM (Fig.1D and Table S4). Similarly, *fpvI* encoding the sigma factor and master regulator of iron homeostasis displayed genetic variations in all DK02, DK06 and DK01 isolates. Interestingly, in intermediate and late isolates of DK19, mutations were observed in the *pvdS* regulator gene, which belongs to the same regulatory network as *fpvI* (Fig.1D and Table S4). The *bfmR* regulator (involved in biofilm maturation, Rhl quorum-sensing (QS) system, and active in acute infections), and the *algU* regulator (involved in alginate biosynthesis), show a certain degree of convergent evolution being mutated in more than half of the isolates. The anti-sigma factor, *mucA*, which modulates the activity of *algU* and causes a mucoid phenotype displayed by several Pa isolates, showed time-dependent frame-shift mutations in all late isolates (Table S5). Several additional regulators such as *mvfR, lasR, fliA, exsA, cdpR, vqsR, rhlR, rpoN* and *sphR* showed lineage-dependent convergence confirming strong evolutionary contingency between lineages (Fig. 1D and Table S4).

In summary, convergent evolution was observed for three categories of master regulators controlling envelope remodeling (*mucA*-*algU*), iron metabolism (*fpvI, pvdS*), and quorum sensing-virulence modulation (*lasR, rhlR, vqsM, mexT, bfmR*) (Fig.1D). Our analysis suggests that common selective forces drive the acquisition of mutations in selected regulatory networks in a patient-independent manner. In addition, evolutionary contingency selects for lineage-dependent variants favoring adaptation to the patients.

### Transcriptional convergence of Pa lineages

We previously suggested that acquisition of several regulatory mutations converted DK02 into a lineage highly adapted to the human airways (5). Limited transcriptional changes were, indeed, observed upon acquisition of mutations affecting the envelope (*algU*), catabolism (*rpoN*) and QS (*lasR-rhlR*), even after three decades of infection. To investigate the transcriptional impact of the lineage-specific and shared mutations identified in the DK02 lineage, we performed RNAseq analysis under conditions mimicking the metabolic conditions in CF (e.g., SCFM).

Pearson correlation analysis applied to the expression data (normalized reads) showed that except for DK02 isolates, the transcriptional correlation among the samples was dependent on the *time of evolution* (early/intermediate ➔ late transition), rather than specific for each *lineage* (Fig. 2). As previously demonstrated (3), all DK02 isolates showed a strong correlation coefficient (Pearson’s correlation coefficient >0.95), independent of time of evolution. Moreover, one late isolate from each lineage clustered with the DK02 transcriptomes suggesting convergent evolution at late times after the onset of chronic infection (Fig. 2A). In contrast, the early isolates of DK19 clustered as a very distinctive class separated from all the samples indicating a very distinctive transcriptional profile at early starting points (Fig. 2A). A similar result is obtained when comparing transcriptional profiles using PCA. All late Pa strains cluster closer to the DK02 strains rather than their specific early strains indicating convergent evolution (Fig. S2A).

**Figure 2.**
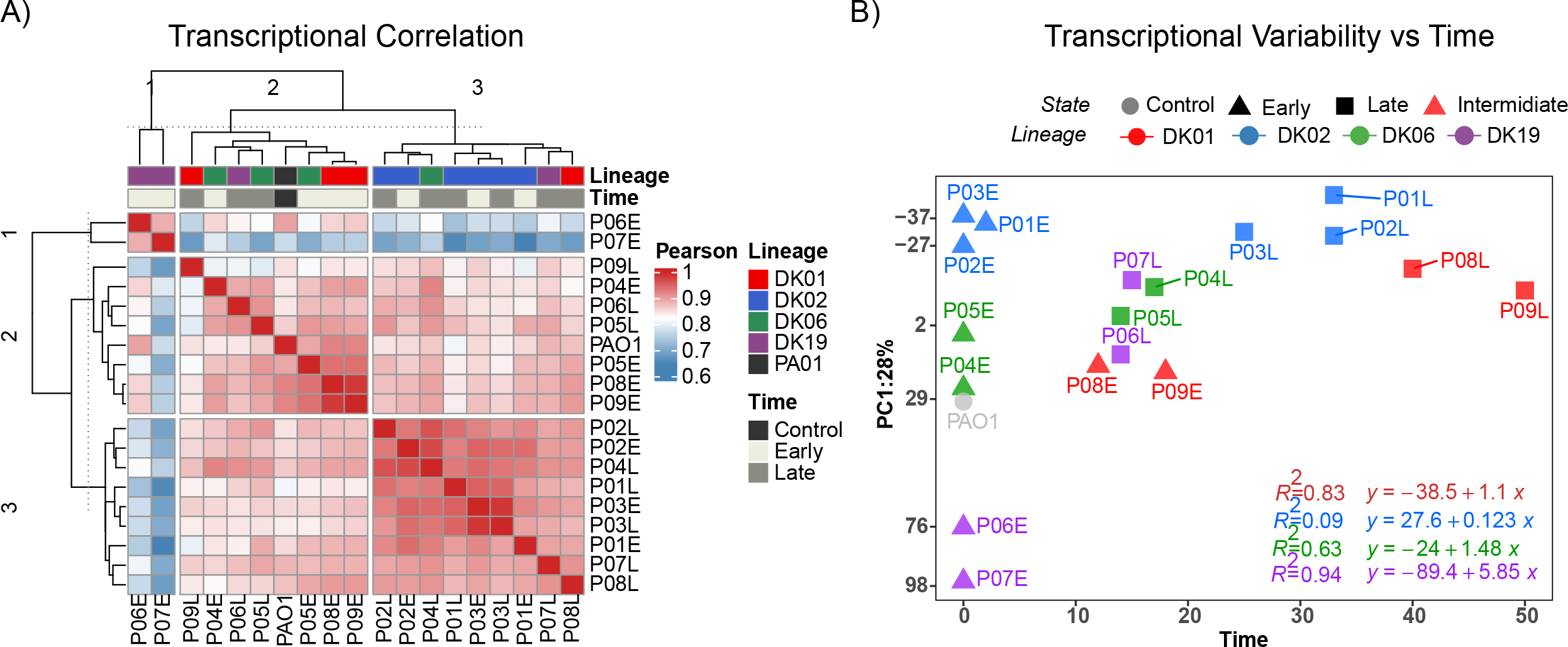
Transcriptional variability over time. A) Transcriptional-based correlation expressed as Pearson’s correlation coefficient (*r*) and visualized as a heatmap of transcriptional profiles. B) Unsupervised Principal component analysis (PC1) loadings for each sample graphed as a function of time. PC1 represents 39% of the explained variability. Lineages dominating each patient are color-coded (DK01 red, DK02 blue, DK06, green and DK19 purple).

Since the time of evolution (years after the chronic infection was diagnosed) seems to play a major role in shaping the lineage phenotype, we compared the transcriptional variability to the length of infection by performing principal component analyses (PCA) on the normalized reads versus time (Fig. 2B). The more evident signature was for DK19, for which the transcriptional variability over time (slope) was 5-to 6-fold higher than that of the other lineages -followed by DK06 and DK01, with an R square of 0.95, 0.63 and 0.83, respectively (Fig. 2B). DK02 displayed an R square and slope close to zero, representing essentially no transcriptional changes during the lineage evolution (Fig. 2B). In contrast, DK01, DK06 and DK19 seem to reflect phenotypic transition states directed towards the stable transcriptional state observed for DK02 (Fig. 2B).

To illustrate the impact of the specific regulatory mutations on the transcriptional network of the DK02 lineage, we included two PAO1 derivative mutant strains harboring the same mutations as the late DK02 isolates. Specifically strain “RegMut” harbors alterations in *mucA-algT-rpoN*, while strain “RegMutΔlasR” harbors an additional deletion in *lasR* regulator (5). Both PCA and Pearson correlation analysis showed that such strains represent late transcriptional states close to those of the DK02 isolates (Fig. S2A, B), and suggesting that the transcriptional state and stability seen already in early isolates of DK02 represents an adaptive maximum, which many/all Pa strains attain over time after infection of the human airways.

### Pathway selectivity at late time points

To further characterize the transcriptional changes associated with the late chronic adaptation states, we identified differentially expressed genes (DEGs) in early/intermediate versus late strains and performed enriched analysis based on KEGG/GO pathway/function classifications. This provided an identification of selective outcomes from the fixed mutations and characterization of their influence on the directionality of the transcriptional program. KEGG and GO enrichment analysis showed that during the adaptive processes, different lineages displayed similar enriched pathways and thus, evolutionary convergence (Fig. 3A, Fig. S4). The lineages converged at late time with higher frequency in 3 major features: i) increase in expression of ABC transporters for sulfur metabolism, ii) decrease in expression of QS regulators controlling e.g., cyano-amino acid metabolism and phenazine biosynthesis, iii) decrease in expression of certain biofilm biosynthetic genes, iv) activation of transcriptional factors related to siderophore uptake/activation (Fig. 3A, Fig. S4).

**Figure 3.**
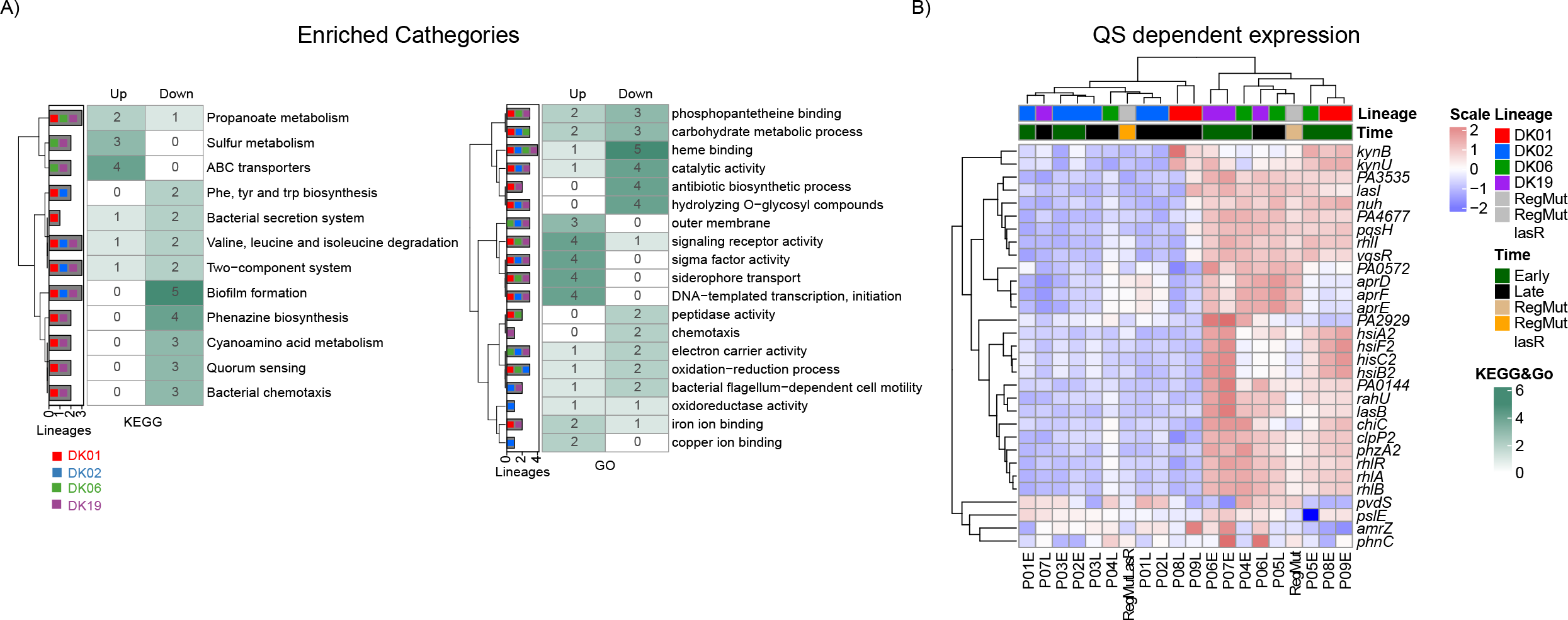
Convergent evolution on different lineages. A) KEGG and GO enrichment counts of the early-late stage are represented as a heatmap of the sum classes. B) Transcriptional patterns of differentially expressed genes of *lasR* regulon. Expressed as log-regularized read counts, scaled for each row, and visualised as a heatmap. Each column represents an analysed sample and is clustered based on the result of pvclust. Lineages dominating each patient are color-coded (DK01 red, DK02 blue, DK06, green and DK19 purple).

Most of these time-associated changes are stably conserved in the DK02 lineage, or present a time dependent regulation (e.g., biofilm downregulation), once more suggesting that all isolates of the DK02 lineage are fully adapted at an early infection stage (Fig.S4). Importantly, our data further suggest that the enrichment of certain mutations at later infection stages may be related to pathway selectivity, showing cases of convergent genomic evolution among different lineages. Furthermore, depending on the evolutionary state of the specific isolate, our data highlight the strong selective pressure for the modulation of QS, as many of the convergent changes are regulated by it (e.g., biofilm, siderophore, phenazine, etc) (Fig. 3B).

Comparing the transcriptional levels of genes regulated by QS in early and late isolates suggests that QS exerts strong negative selectivity on late isolates, both in a *lasR-rhlR* mutation-dependent (e.g., P01L and P08L) and -independent manner (e.g., P09L)(Fig. 3B) (15). The downregulation of QS may indicate selection for loss of function mutations for the entire pathway, or modulation of pathway expression. To distinguish between these possibilities and to further investigate the changes in the excretion of other important chemical compounds, we performed an exo-metabolomic analysis of the secreted molecules by these lineages.

### Metabolic distinctions during different evolutionary stages

Metabolomic profiles from stationary phase cultures were analyzed by means of liquid chromatography coupled to MS (LC-MS). The molecular masses obtained were aligned and quantified for the different isolates (Table S6). Unsupervised PCA profiling of the total exo-metabolomes clearly documents different outcomes from early (cluster A) and late isolates (cluster B-C), except for DK02, which once more shows no variation between the different time points of the isolates (cluster B) (Fig. 4A). Clusters C and B show reduced or undetectable levels of oxo-C12-HSL, C4-HSL and HHQ, with a concomitant reduction in phenazine, pyocyanin and rhamnolipids (Fig. 4C). Cluster A represents bacteria, like PAO1, with normal levels of oxo-C12-HSL, HHQ, phenazine, pyocyanin and rhamnolipids (Fig. 4C).

**Figure 4.**
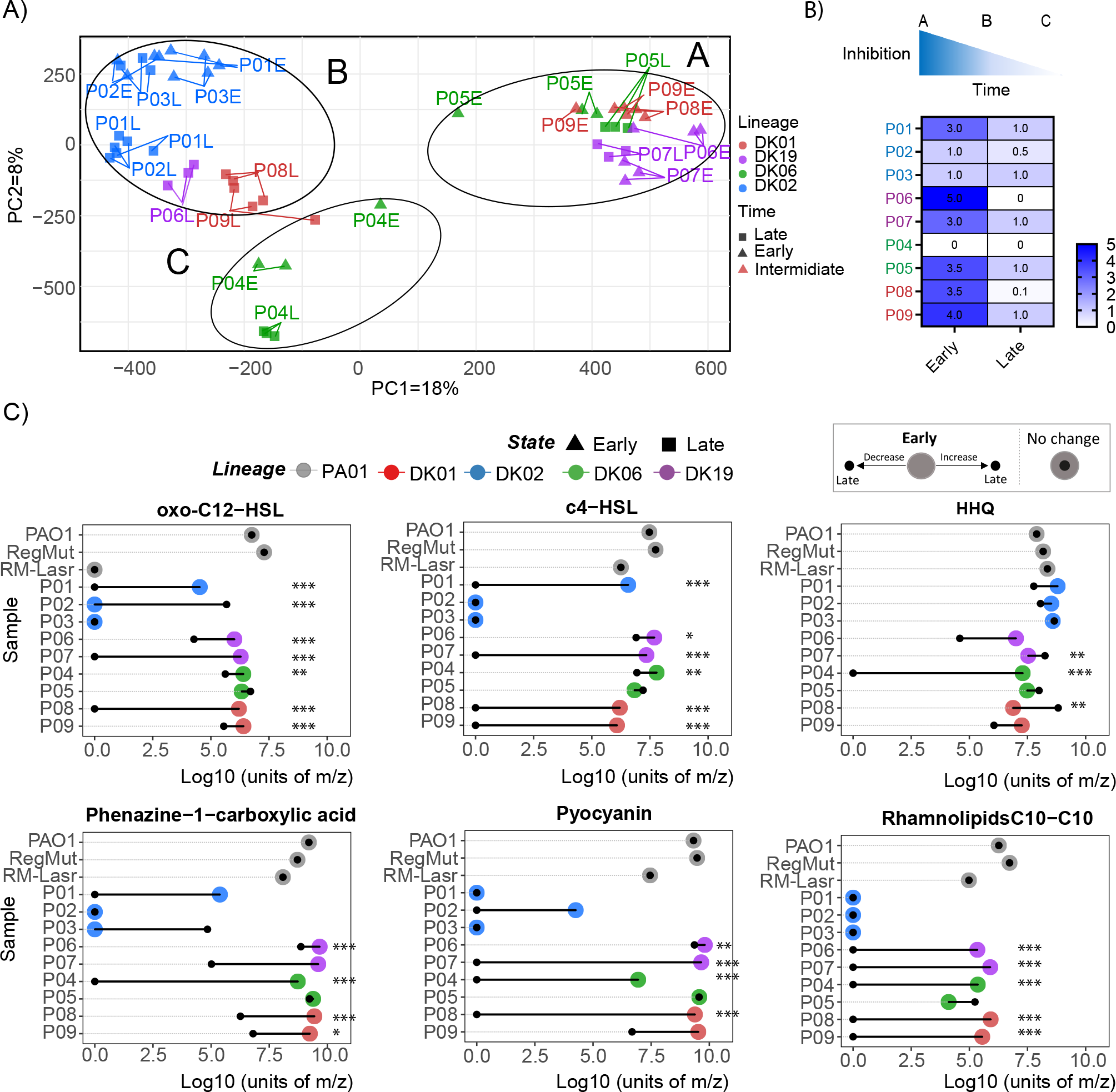
Metabolomic distribution in the different samples. A) Unsupervised principal component (PCA) analysis performed on the total exo-metabolites. Lineages are colour-coded. B) Inhibitory effect of secreted supernatant tested on sensitive bacteria (*B. subtilis*). C) Quantification of relevant metabolite, QS regulators oxo-C12-Homoserine-lactone (oxo-C12-HSL), N-butanoyl-L-Homoserine-lactone (C4-HSL), 2-heptylquinolin-4(1H)-one (HHQ), Phenazine 6-carboxylic acid, Pyocyanin and C10-C10 Rhamnolipid. Secreted metabolites from Early samples are represented as colour circles and the related Late with a dark point. Samples coming from the same patient are connected with a line. Lineages dominating each patient are color-coded (DK01 red, DK02 blue, DK06, green and DK19 purple).

Most isolates showed only small variations in HHQ production, although statistically significant reductions were observed in one DK06 late isolate (Fig. 4C). Interestingly, the genes controlled by C4-HSL were downregulated in the late isolates (Fig.3B), suggesting that the production of this QS molecule does not impact the QS network downstream, further suggesting that mutations in the central QS regulatory genes (*lasR-rhlR*) might govern i) the transcriptional reduction of the network regulon genes, and ii) the modulation of the QS secreted molecules.

As expected, the PAO1 *RegMut*Δ*lasR* strain displayed a complete depletion of oxo-C12-HSL production (Fig. 4C). Moreover, it displayed a metabolome profile in between clusters A and B, with 1-2 Log_10_ reduction (200 times) in C4-HSL, rhamnolipids, pyocyanin and phenazines. However, this reduction was not as drastic as in many strains from cluster B, where the levels of these molecules were non-detectable (Fig. 4C).

Since QS is associated with virulence, we tested the inhibitory properties of these secreted molecules against sensitive bacteria, and it was clear that early strains showed the highest levels of virulence (cluster A). Moreover, in all cases, the late, evolved isolates showed reduced virulence, in line with the genomic and transcriptomic information (clusters B and C) (Fig. 4B).

There was a high prevalence of some QS-regulated molecules, but only small variations in the production of HQN, HNQO and other quinolones from different lineages at different time points (Fig. S4). All strains showed stable excretion of the siderophore pyochelin, whereas the excretion of pyoverdine was reduced in some late isolates. Increased excretion of a molecule, identified as Norleucine, was seen in the late isolates, whereas reduced excretion was observed in the early isolates compared with DK02 (Fig. S5).

In conclusion, the metabolomics data suggest that adaptation of Pa to human airways entails a strong selective pressure for loss of virulence-associated molecules (e.g., phenazines, pyocyanin and rhamnolipids) and a maintenance of several important eco-physiological properties (e.g., production of siderophores and quinolones). The loss of production of virulence-related molecules and modulation of the production of physiologically important molecules further suggest different functional roles for these families of molecules and, consequently, stringent modulation of their QS-mediated regulatory components.

## Discussion

We and others have previously documented that bacterial colonization and adaptation in the airways of CF patients constitutes an interesting model for studying microbial evolution in complex, dynamic environments (4, 11, 14). With a special focus on the environmental bacterium *P. aeruginosa* (Pa), the process of migration from the environment to human airways has been monitored by genotyping and phenotyping various collections of Pa isolates from pwCF. It has been convincingly documented that after years of bacterial colonization, Pa isolates derived from the patients have gone through extensive genetic and phenotypic alterations, which eventually result in the conversion of the Pa generalist type of organism to one which behaves much more like a niche specialist (11).

Here we follow 4 different lineages from pairs of isolates independently evolving in the lungs of different patients for periods corresponding to 35,000-150,000 bacterial generations. In line with previous results from our lab, we see accumulation of mutations since the persistent infection was established (Fig. 1C and Fig. S3B). Perhaps the most important consequence of these mutations is a significant change in the genome-wide transcriptional profile for all the lineages (except for DK02). Although the strains acquire mutations at similar rates, the functional roles of these mutations may differ depending on the state of adaptation of each specific isolate and on the mutations fixed at earlier stages of the infection. Our data further suggest that when a lineage has reached an adaptation maximum, additional fixation of mutations in regulatory genes (including additional transcriptional changes) may entail fitness costs for the population, and therefore such mutated isolates will be diluted out in the population. Notably, limited genomic convergence, shown as gradual accumulation of mutations during the evolution of the lineages, resulted in high convergence of the transcriptional programs. Due to the plasticity and functional redundancy of Pa genome, we often see that many genetic routes have similar phenotypic impact, i.e., similar global transcriptional network. Moreover, mutations that are fixed from early stages of colonization may direct the order of selection of new mutations in different arrays of genes (11). This illustrates cases of evolutionary contingency or epistatic interactions necessary to cope with the continuous changing environment.

The transcriptional convergence program associated with the time of infection occurred in a patient-independent manner, suggesting that similar selective pressure dominate the human airway environments at late time points. Convergence of the transcription and regulatory mutational profiles was illustrated by the gain of mutations in 19 well-known pathoadaptive loci from Pa, mutated across the different analyzed lineages (11) (Fig. S8C, Table S5). Moreover, mutated genes could be grouped into functional categories such as envelope modifications, catabolic modulation, biofilm regulation, changes on iron response, and antibiotics resistant genes (Fig. S1D, Fig. S4). When a Pa lineage has entered the airways and developed into persistent infection, there is an array of functional mutations that need to be introduced in the genome, so the population acquires a steady state of transcriptional variation, like that seen for DK02.

One key feature that clearly distinguishes between early and late isolates is the down-regulation of QS. In the investigated isolates, reduction in QS was associated with mutations in the major QS modulators *lasR-rhlR* (e.g., DK02, DK19 and DK01) genes in most of the late isolates, or through mutations in the virulence modulator *gacS/retS* (e.g., *ladS* and *pprA* for DK06). In the DK19 lineage, one isolate had a deletion in *lasR* with drastic effects on the transcriptional program, converting the transcriptome to that described for the DK02 lineage (Fig. 1D and Fig. S6). But, the *lasR-rhlR* mutation alone could not explain the drastic reduction of HSL and virulence related moieties observed in late isolates (e.g., Phenazines), as shown for the PAO1 RegMutΔlasR strain with a *lasR* deletion, which shows a reduction of QS, less drastic than what is observed in the late isolates. It is therefore likely that the mutational profiles acquired during persistent infection may comprise loss of function mutations within the regulatory networks implicated in the modulation of QS.

QS down-regulation was observed mainly in isolates, which were obtained many years after infection of the patient airways (e.g., 1-3 decades). This may suggest that at early times, QS may be important for the establishment of the infection. In fact, reduction of QS is usually associated with increased probability of persistent infection (15). During the progress of the infection in time, the CF-airways show biotic and abiotic physiological variations. Usually, there is a dysregulated immune system (e.g., increase the population of neutrophils and immune cells), and changes in the biotic environment in the lung, associated with decreases in microbiological diversity and dominance of one or a few opportunistic bacteria (1). Moreover, expression of virulence factors and QS molecules may be energetically costly and due to fitness pressure, loss of function mutations are acquired when the population diversity declines. Therefore, it is possible that the evolutionary convergence observed in the isolates could be related to a similar selective force governing the CF-lung environment at a late time as a response to an eco-physiological variation (16).

Although the production of AHL and the transcriptional network modulated by AHLs was eliminated in most of the late isolates, some QS-regulated molecules were synthesized, probably independently of the QS network. Among the molecules produced by the Pa isolates at late time points, siderophores and quinolones could be associated with specific functional roles of these molecules. For example, siderophores, commonly known as metal chelators implicated in iron and other metals’ homeostasis, were produced by all the isolates, at least one type (e.g., pyoverdine vs pyochelin). Moreover, HQNO and other AQNO quinolones are redox molecules implicated in the modulation of the immune response and virulence factors for other bacterial warfare (17). It is possible that the selective value of AQNOs is related to immune modulation more than to virulence, as the more virulent-associated molecules Phenazines/Pyocyanin and rhamnolipids (18) showed a drastic decrease with time. In summary, both iron homeostasis modulation and probably immune modulation may be key features for the adaptation of Pa.

Finally, associating evolutionary data with patient information could be used as a proxy for the development of biomarkers to determine the patient’s prognosis and/or disease development. We believe that evolutionary studies like the one presented here could help pinpoint genomic determinants associated with pathway-specific selectivity, providing an easier genomics-phenotypic association. Moreover, it could provide a proper biomarker of the infection stages and improve treatment options for the patients.

## Materials and Methods

### *P. aeruginosa* CF isolate collection, ethics approval and consent to participate

Clinical isolates were obtained from sputum samples from 9 patients with cystic fibrosis attending or that have attended the Copenhagen Cystic Fibrosis Center at University Hospital Rigshospitalet, Copenhagen, Denmark. Sputum sampling was part of routine clinical visits in the CF clinic and not performed for the purposes or intent of this research. The use of the isolates was approved by the local ethics committee of the Capital Region of Denmark (Region Hovedstaden; registration numbers ^H-21078844^).

Isolation and identification of *P. aeruginosa* from sputum was carried out as previously described (19). The *P. aeruginosa* collection included pair of isolates from each of the patients, one taken at the beginning of the chronic infection, and one taken after a period of 15 to 40 years depending on the patient/lineage. Time in which pair of isolates were collected is summarized in Table S1.

### Laboratory bacterial strains

*P. aeruginosa* reference strain PAO1 was used in this study together with two isogenic mutants (Regmut and RegmutΔlasR) previously constructed in the lab associated to DK02 evolutionary history (5). Regmut, consisted of a triple mutant based on specific *mucA-, algT- and rpoN* alterations, and RegmutΔlasR included an extra deletion of *lasR* gene, giving a quadruple mutant configuration.

### Comparative genomics

Genomic DNA was extracted and purified from overnight liquid cultures of bacterial single colonices using a DNeasy Blood and Tissue kit (Qiagen). Genomic DNA libraries were prepared using a Nextera XT DNA Library Prep kit (Illumina), and libraries were sequenced on either a MiSeq (69 libraries) or NextSeq 500 platforms (84 libraries), generating 250 or 150 base paired-end sequencing reads, respectively. Sequencing reads were trimmed, and low-quality reads and potential contamination from adapters were removed using Trimmomatic (v 0.35) tool (20). Reads were mapped against the *P. aeruginosa PAO1* genome (NCBI: NC_002516.2) using BWA-MEM algorithm (21), and duplicated reads were marked using Picard tools. Genome Analysis Toolkit (GATK) (22) was used to re-align and call SNPs. SNPs were extracted if they met the following criteria: a quality score of at least 50, a root-mean-square (RMS) mapping quality of at least 25 and a minimum of three reads covering the position. Microindels were filtered based on a quality score of at least 500, an RMS mapping quality of at least 25 and support from at least one-fifth of the covering reads. Variations unique to each clone belonging to the same lineage were used to determine potential transmissions and to estimate an average evolutionary distance.

### Phylogeny reconstruction of CF isolates

Evolutionary analyses were conducted in MEGA11 (23). For this purpose, concatenated sequences of only the SNPs of the 19 CF isolates were aligned to the positions of the nucleotides in the genome of the reference strain PAO1. There were a total of 76662 positions in the final dataset. Evolutionary history was inferred by using the Maximum Likelihood method and General Time Reversible model (23) and 500 bootstrap was set for analysis confidence. Boostrap tree is shown.

### Estimation of bacterial evolution rates

Evolution rates were assessed as the number of SNPs per genome size per generation as previously (4). Generation times, assembly size, and estimation of bacterial evolution is summarized in Table S2. For SNPs, the number of missense, stop and synonymous mutations accumulated between the pairs of early and late isolate was used. For estimation of bacterial generations, we calculated the growth rate of each isolate in SCFM media which gave us an average doubling time of 140.04 ± 49 min. This value was already on the range of previous published data from *in vivo* doubling time ratios (24). The number of bacterial generations elapsed over time was calculated as the sum of generations from the year of isolation of the early to the late. As a proxy of genome size for each lineage we assembled the genomes of one single pair of isolates: for DK01(P09I-P09L), DK02 (P02E-P02L), DK06 (P04E-P04L) and DK19 (P07E-P07L). For this, pair end reads were assembled into contigs using Spades (25) and quality was evaluated with QUAST (26).

### Library preparation and RNA sequencing

Single colony cultures were grown in SCFM media (inoculation with OD_600_□=□0.05) at 37 °C under shaking conditions (200□rpm) to mid-exponential phase (OD_600_□=□0.35-0.5). RNA was extracted with RNeasy Mini Kit (Qiagen) according to the manufacturer’s instructions. Transcription was blocked applying RNA Protect Bacteria solution (Qiagen). RNA was quality-checked using an Agilent Bioanalyzer 2100 (Agilent Technologies) (RIN > 9). For all other samples used in this study, 100□ng of total RNA was used as input for the generation of RNA libraries with Illumina® Stranded Total RNA Prep, Ligation with Ribo-Zero Plus kit and following manufacture’
ss instructions. After quality and size distribution check on DNA HS chips on an Agilent Bioanalyzer 2100 machine, libraries were pooled in equimolar amounts and sequenced on an Illumina NextSeq 500 machine. An average of 10-15 million reads per samples with 2×75-bp-long reads per sample. Mapping was performed using the PA14 genome as a reference.

### Comparative transcriptomics

Reads were trimmed, and low-quality reads and potential contamination from adapters were removed using Trimmomatic (v 0.35) tool (20). Reads were further processed using SortMeRNA tool (v 2.1) (27) to remove reads generated from residual rRNA transcripts. As DK19 is the PA14 lineage, and to have a better read alignment, reads were mapped against the *P. aeruginosa UCBPP-PA14* genome (NCBI: NC_008463.1) using BWA-MEM algorithm, and duplicated reads were marked using Picard tools. Reads mapping on each annotated coding sequence were counted using htseq-count version 0.7.2 (28) imported and processed in Rstudio (29).

### Transcriptional analysis

Counts were normalized using log_2_ negative binomial transformation performed using the rld Transformation function contained in the R package DESeq2 (30) with option blind set as “True”. Normalized counts were used to evaluate whole transcriptome similarities using hierarchical clustering analysis (HCA), principal component analysis (PCA) and k-mean clustering on selected normalized data. HCA was performed using the function “pheatmap” in the R package complexheatmap (31). Pearson’s correlation coefficient (cor()) was applied on the normalized data as a distance method. Principal component analysis on normalized reads counts was performed using prcomp() function with scale option set as “FALSE”. Differential gene expression (DEG) analysis between transcriptomes was performed using the R package DESeq2, considering statistically significant genes with a Log_2_(FoldChange)□≥ □|2| and an adjusted p value ≤ 0.01 (30). DEGs were inspected and functional class enrichment was performed using the provided the R package ClusterProfiler (32) for KEGG and GO categories with default settings. The convergent enriched pathways, similar up/down regulation for each pair of samples, were evaluated by quantifying the frequency that that enriched categories appear in at least 2 pair of samples.

### Comparative metabolomics

Single colonies were isolated on LB agar plates and transferred to a tube containing 2 mL SCFM+GlcNAg medium. This pre-culture was normalised to OD:0.05 using fresh medium and added to a flask. Samples were incubated at optimal temperature/shaking conditions. Bacterial cultures were isolated to stationary phase when no OD variations were observed, OD_600_ = 2-3, t = 24 h for early strains and 36 h for late isolates. For all the isolates, 2 ml of actively growing bacteria were centrifuged (6000 g, 10 min), and the supernatant and pellet were used for further experimentation. The supernatant was 0.22µm filtered and concentrated on a speed vacuum R.T.. Dried pellets were stored at -80 °C. The pellets were concentrated 10 times and resuspended 50:50 MQ water: Methanol. The samples were run on a Vanquish Duo UHPLC binary system (Thermo Fisher Scientific) coupled to IDX-Orbitrap Mass Spectrometer (Thermo Fisher Scientific, USA). The compounds separation was achieved in reverse phased using a Waters ACQUITY BEH C18 (10 cm × 2.1 mm, 1.7 μm) column equipped with an ACQUITY BEH C18 guard column kept at 40 °C and mobile phase consisting of MilliQ water + 0.1% Formic acid (A) and Acetonitrile + 0.1% Formic acid (B) at a flow rate of 0.35 mL/min as previously described (33). The MS acquisition was set in positive-heated electrospray ionization (HESI) mode with a voltage of 3500 V acquiring in full MS/MS spectra (Data dependent Acquisition-driven MS/MS) in the mass range of 70–1000 Da. The DDA acquisition settings were as follows: automatic gain control (AGC) target value set at 4e5 for the full MS and 5e4 for the MS/MS spectral acquisition, the mass resolution was set to 120,000 for full scan MS and 60,000 for MS/MS events. Precursor ions were fragmented by stepped High-energy collision dissociation (HCD) using collision energies of 20, 40, and 60. All the analyses were carried out in biological triplicates. LC-MS chromatograms were aligned and quantified using Mz-mine with default parameters (34). Masses were further process in R. Missing values were given a 0 value. For unsupervised clustering, acquired masses were normalized in R with a negative binomial normalization, applying a variant stabilizing normalization inside the DeSeq2 package (30), with option blind set as “True”. Molecular masses were confirmed by comparing retention time to commercial standards (e.g., oxo-C12-HSL, phenazines, pyocyanine and rhamnolipids), development of pathways synthetic mutants (e.g., deletion *pqsABC* and *pvdI* quinolones and pyoverdine respectively) and comparing to virtual libraries (e.g., Rhamnolipids) (34). Masses were validated by analyzing their fragmentation profile to that previously stored in GNPS library (35). Statistical analyses were performed with unpaired Log10 multi-t-test inside GraphPad. Differences were considered statistically significant at p < 0. 05.

### Inhibitory assay

A pre-culture of *Bacillus subtilis* was grown ON and normalized to OD0.01. 200 µl of cultures were spread to LB-agar. The plates were dried and whatman paper discs were attached. Then, 5µl of the concentrated bacteria filtered and concentrated 10 supernatant were added to the dics. Plates were incubated at 37 O.N. As positive control 1500 µg of Ciproflaxin was added (5 µl at 300 µg/ml) and a negative control fresh concentrated SCFM medium.

### Data Availability Statement

Whole genome sequences of isolates P01E, P01L and P03E were downloaded from previous published data from the Sequence Read Archive (SRA) study ERP002277. The same applies for isolates P05E and P06E, which were obtained from the SRA study with accession number ERP004853. Raw sequences of the rest of the WGS and RNA-seq have been deposited in the SRA under BioProject ID PRJNA991306. See Table S1 for the accession codes of individual isolates data.

## Author contributions

A.E., C.A.C, S.M and H.K.J. conceived and designed the study. A.E. and C.A.C performed the genomic and transcriptomic experiments. A.E. and D.R. performed and analyzed the metabolomics experiments. A.E. and C.A.C analysed the data and all authors contributed to the interpretation of the data. A.E., C.A.C, H.K.J, S.M. and R.L.R wrote the draft manuscript, and all authors edited the paper.

## Acknowledgements

We thank Elio Rossi, Oihane Irazoqui and Maria Pals Bendixen for their insightful comments and discussions. We also thank the Sequencing and Analytics departments in the Novo Nordisk Center for Biosustainability biofoundry for helping with running RNAseq and metabolomic samples. This work was supported by a Novo Nordisk Foundation Challenge Grant NNF19OC0056411 to H.K.J.

## Figure Legends

**Figure supplementary 1. Genes shared and non-shared within each of DK01, DK02, DK06 and DK19 lineages**. Percentage of common or isolate specific mutated genes within early (A) and late (B) isolates is shown. Newly mutated genes between early to late pair of isolates within each lineage are analyzed based on (C) Percentage of genes being common or isolate specific and, (D) Functional categories of common mutated genes.

**Figure supplementary 2. Transcriptional variability**. A) Unsupervised Principal component analysis (PC1-PC2) loadings of the different isolates. B) Transcriptional-based correlation expressed as Pearson’s correlation coefficient (*r*) and visualized as a heatmap of transcriptional profiles from the different isolates.

**Figure supplementary 4. Enriched pathways analysis**. KEGG and GO categories of the differentially expressed up and downregulated genes between the early and late pairs.

**Figure supplementary 3. Genomic and transcriptional changes occurring in DK19**. A) Functional genomic changes occurring during evolutionary adaptation B) Visualization using pathview (XY) of the differentially expressed genes between the late DK19 isolates.

**Figure supplementary 5. Selected metabolites present in the different lineages**. C) Quantification of relevant metabolite, 4-hydroxy-3-nitroquinolin-2(1H)-one (HNQ), 2-heptyl-4-hydroxyquinoline N-oxide (HQNO), norleucine, pyoverdine, pyochelin, aeruginoic acid and alkyl hydroxyquinoline (AQNO).

**Figure supplementary 6. DK02-associated enriched pathways**. A) Transcriptional patterns of differentially expressed genes. Expressed as log-regularized read counts, scaled for each row, and visualised as a heatmap. Each column represents an analysed sample and is clustered based on the result of pvclust. B) Illustration of the activation of arn-pathway.

**Figure supplementary 7. Total and Pathoadaptive gene mutation in the CF isolate collection**. Summary of total number of genes being mutated representing (A) strain specific accumulation or, (B) commonly mutated, between early or late isolates from the four lineages analyzed. C) Non-synonymous and indels mutations found within pathoadaptive genes.

