## Supplementary figures and images for "Adaptive evolution of *P. aeruginosa* in human airways shows phenotypic convergence despite diverse patterns of genomic changes"

### Supplemental Figure 1

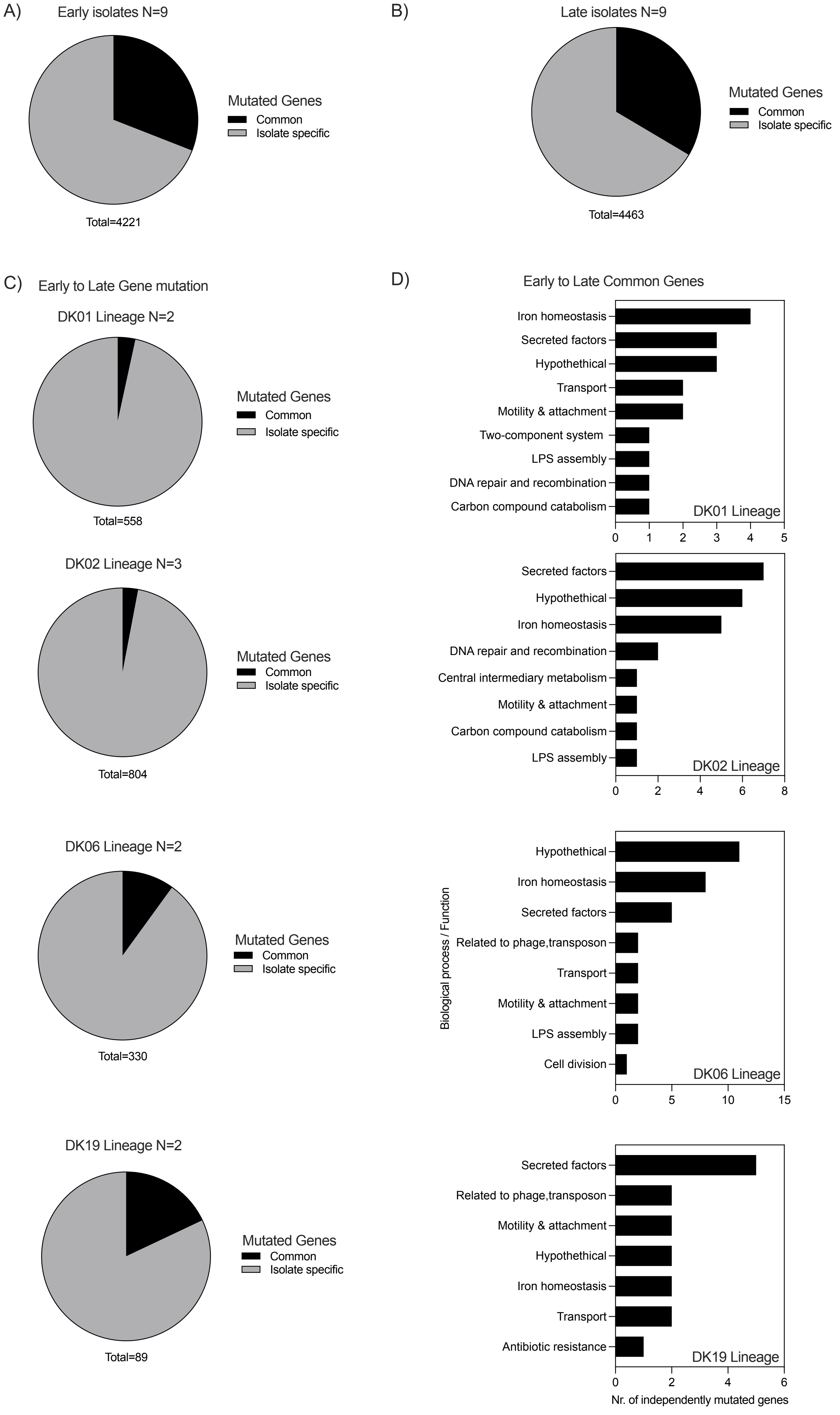

### Supplemental Figure 2

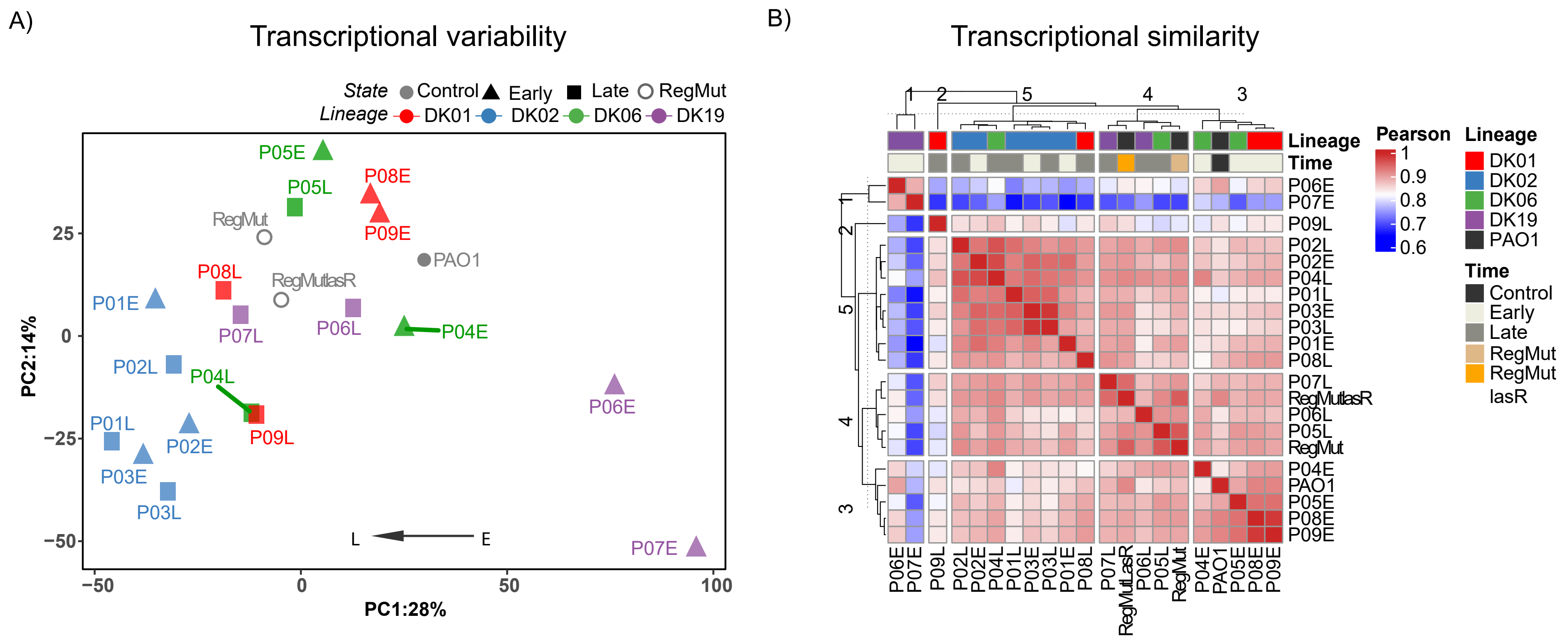

### Supplemental Figure 3

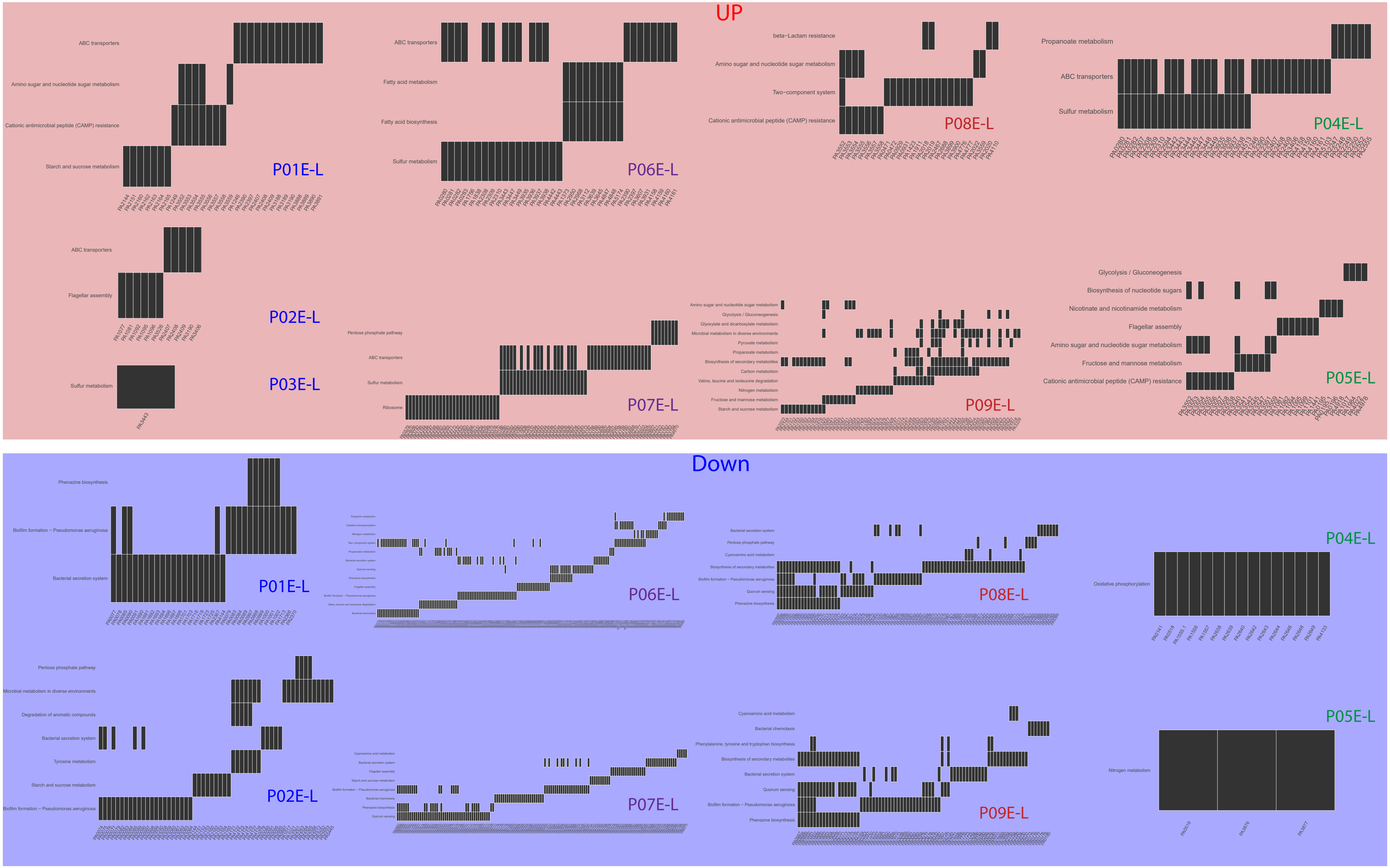

### Supplemental Figure 4

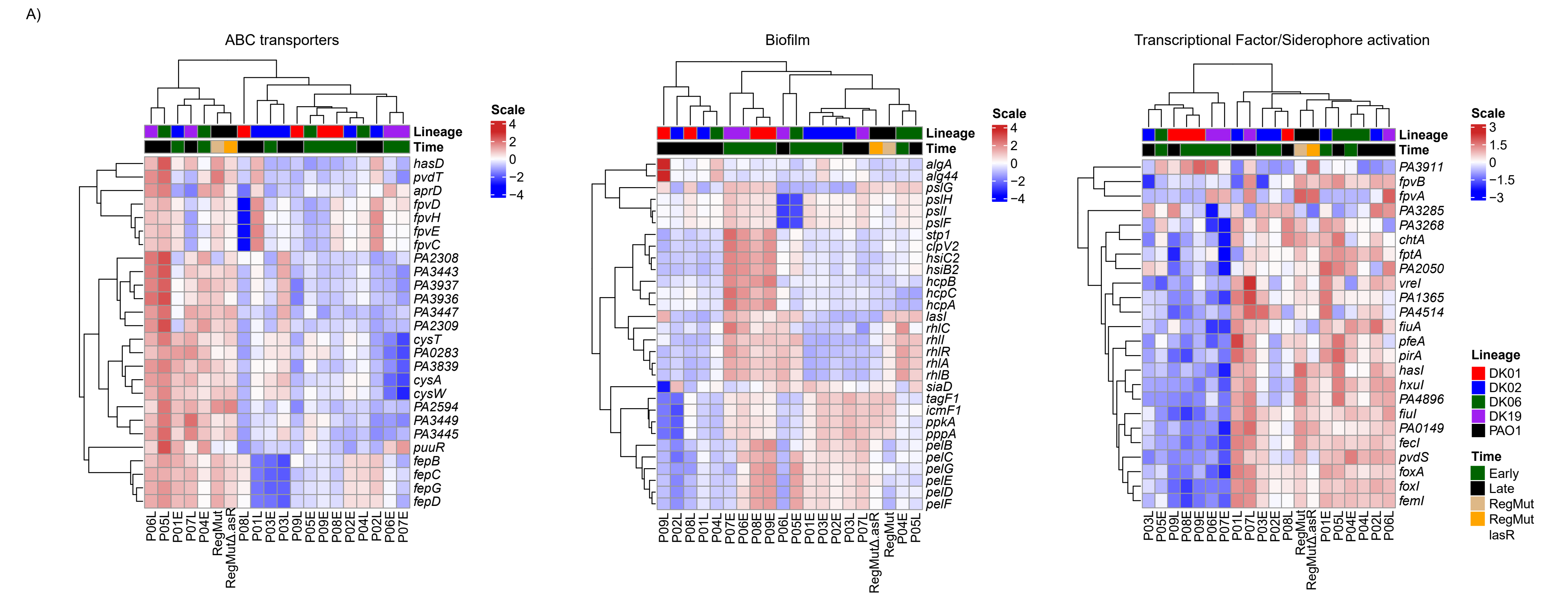

### Supplemental Figure 5

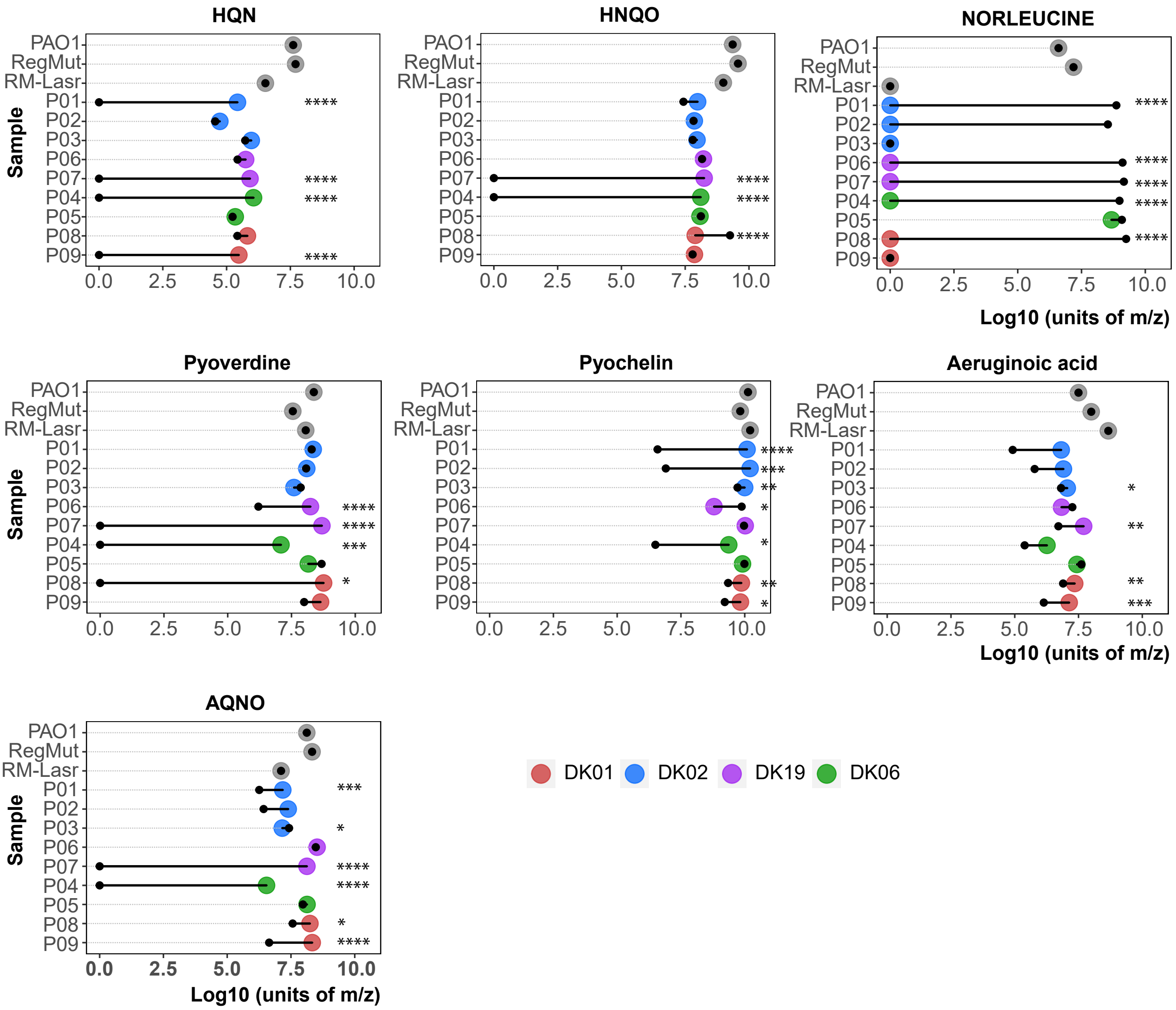

### Supplemental Figure 6

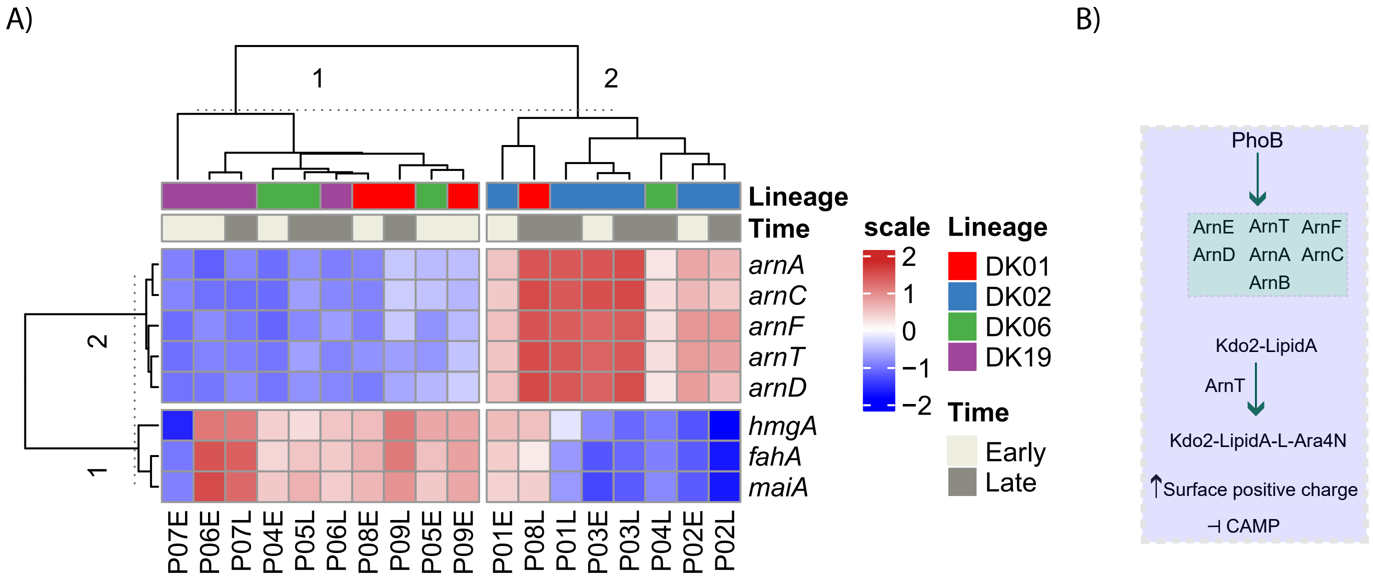

### Supplemental Figure 7

A)

#### Mutation Selectivity

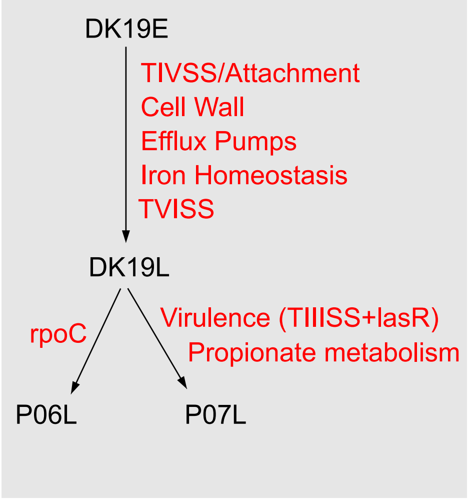


B)

#### P07LvsP06L

Virulence

Metabolism

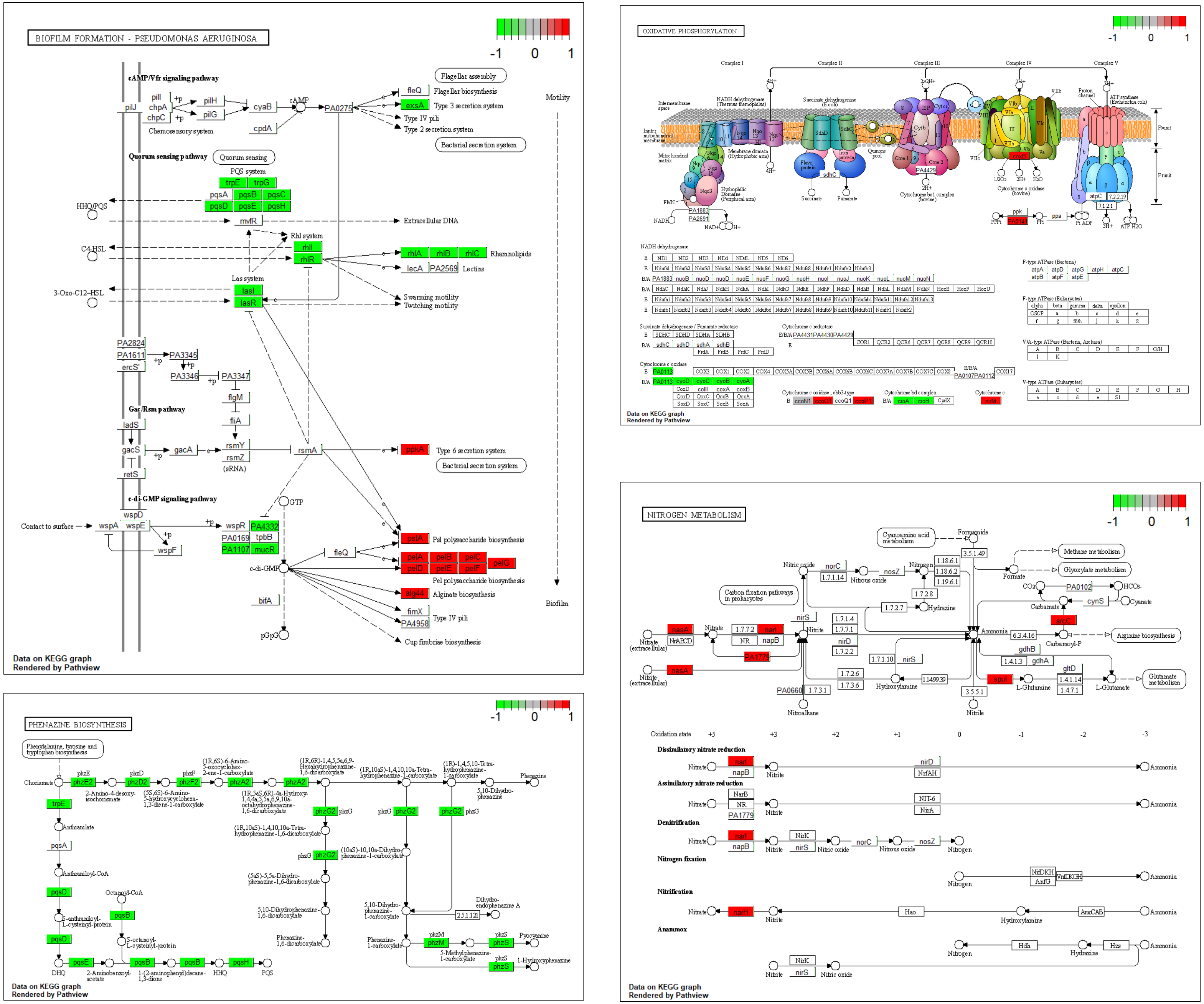
